# Comparative proteomics reveals a conserved core of tegumental proteins in parasitic flatworms

**DOI:** 10.64898/2026.04.22.720116

**Authors:** Inés Guarnaschelli, Analía Lima, Rafael Velazco, Monika Bergmann, Matías Preza, Javier Calvelo, Marcela Cucher, Mara Cecilia Rosenzvit, Klaus Brehm, Andrés Iriarte, Uriel Koziol

**Author notes:** Corresponding authors: Andrés Iriarte: Laboratorio Biología Computacional, Unidad Académica Desarrollo Biotecnológico, Instituto de Higiene, Av. Dr. Alfredo Navarro 3051, CP11600, Montevideo, Uruguay., Uriel Koziol: Sección Biología Celular, Facultad de Ciencias, Iguá 4225, CP11400, Montevideo, Uruguay. int. 144. Unidad de Ingeniería de Proteínas, Institut Pasteur de Montevideo, Montevideo, Uruguay (Analía Lima).

## Abstract

Parasitic flatworms, including cestodes and trematodes, are covered by a specialized syncytial tegument that mediates nutrient uptake and host–parasite interactions. While the tegument of trematodes has been extensively characterized, its molecular composition in cestodes remains largely unknown. In this work, we performed a comparative proteomic analysis of the tegument of three cestode species, including larval and adult stages: *Hymenolepis microstoma*, *Mesocestoides corti* (syn. *M. vogae*) and *Echinococcus multilocularis*. Using stringent enrichment criteria relative to whole-worm extracts, we identified hundreds of tegument-enriched proteins in each species. Comparative analyses revealed a conserved core of tegumental proteins shared among all three species, including members of the Tegument Allergen-Like (TAL) family, vesicular trafficking components and calcium-sensing proteins, and identified candidates for nutrient uptake activities such as glucose and nucleoside transporters. Further comparative analyses revealed a set of shared tegumental proteins with the trematode *Schistosoma mansoni*, including conserved proteins that are specific to parasitic flatworms, supporting the existence of a conserved ancestral tegumental proteome. Finally, we confirmed tegumental expression of several candidate genes in *H. microstoma* and *E. multilocularis*, and demonstrated regionally restricted gene expression among tegumental cytons, suggesting functional specialization within the syncytial tegument. Altogether, these results reveal an evolutionarily conserved composition of the tegument of parasitic flatworms, providing a foundation for future work targeting this critical host–parasite interface.

## Introduction

Parasitic flatworms are causative agents of important diseases in humans and domestic animals [1–3]. Most parasitic flatworms, including cestodes, trematodes and monogeneans belong to the monophyletic group Neodermata [4]. One common characteristic of these parasites is that their bodies are covered by a highly specialized syncytial tegument, also known as neodermis [5]. This tissue is the main contact surface between the parasite and the host (especially in the case of cestodes, which lack a digestive system). As such, it is essential for host-parasite interactions, including nutrient uptake and immunomodulation [6–8]. The tegument consists of a superficial and continuous distal region, which lies on top of a basement membrane, and of nucleated cell bodies (known as cytons) that join the distal region by thin cytoplasmic bridges (Fig. 1A). New proteins are continuously synthesized within the cell bodies and transported to the distal tegument [9–11].

**Fig. 1.**
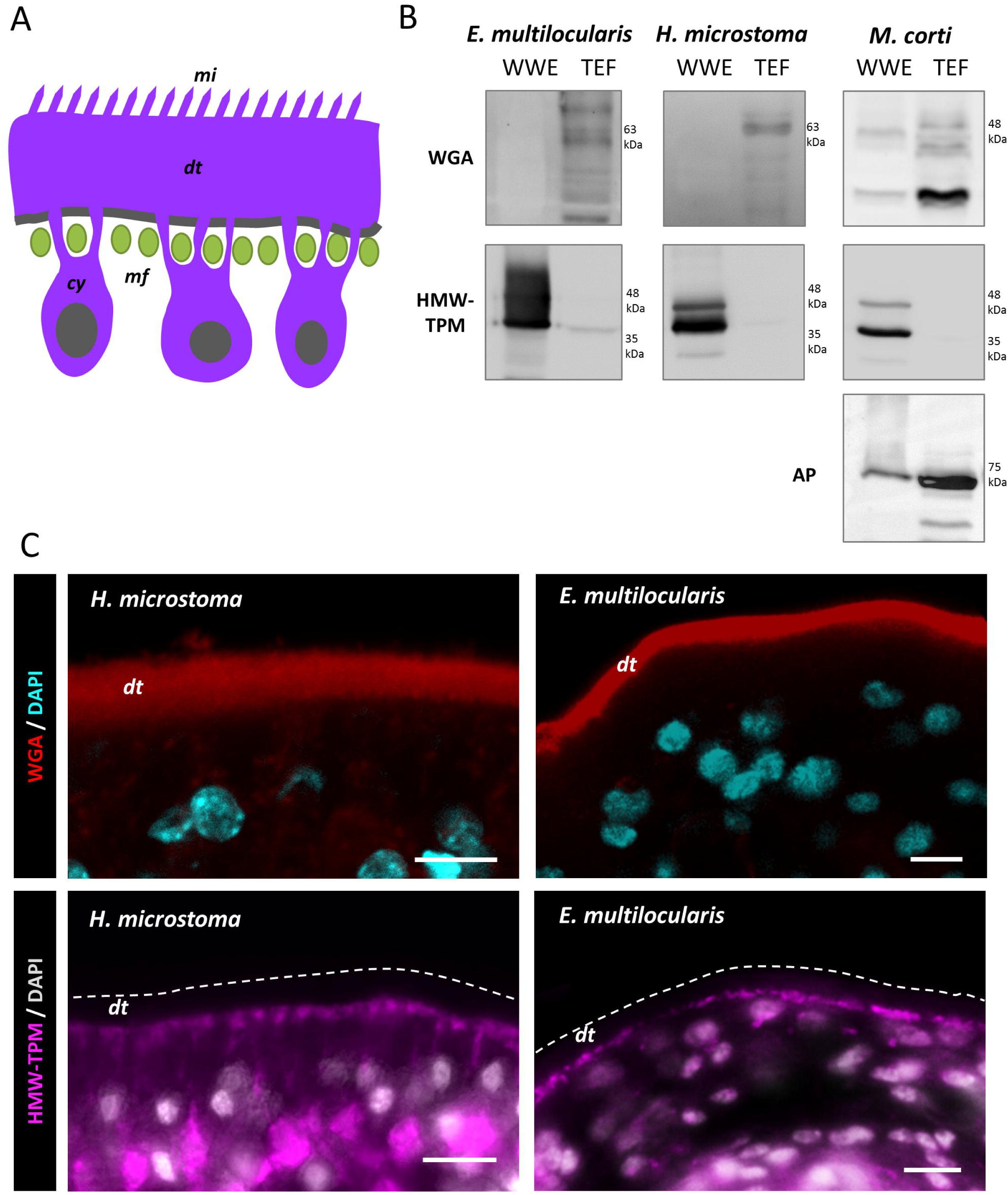
Validation of the tegument isolation method. (A) Schematic drawing of the syncytial tegument of cestodes. (B) Western Blot experiments were conducted on whole worm extracts (WWE) and tegument enriched fractions (TEF). Glycoconjugates detected with Wheat Germ Agglutinin (WGA) are enriched in TEFs with respect to WWE. This is also the case for the enzyme alkaline phosphatase (AP) in *M. corti* TEFs. High molecular weight tropomyosins (HMW-TPMs), which are muscle markers, are depleted in TEFs. (C) Labeling of sections of *H. microstoma* and *E. multilocularis* with fluorescent WGA shows that glycoconjugates are located mainly in the distal tegument. Immunodetection of HMW-TPMs confirms their absence in the distal tegument, and its localization in muscle fibers located immediately below. dt: distal tegument, cy: cytons, mi: microtriches, mf: muscle fibers. Scale bars: 5 µm.

In the case of trematodes, the molecular composition of the distal tegument has been analysed by proteomic methods using highly enriched tegumental fractions. The most extensive results have been obtained in the human pathogen *Schistosoma mansoni* [12], but similar results were also observed in other trematodes such as *Fasciola hepatica* and *Opisthorchis* spp. [13–15]. These studies have identified important enzymes, transporters, structural proteins and antigens in the tegument of trematodes, and experimental vaccines using recombinant forms of tegumental proteins are currently undergoing clinical trials [16–18]. In addition, single cell RNA sequencing (scRNA-Seq) of the trematode *S. mansoni* has shed light on the cellular heterogeneity of tegumental cytons and of their progenitors, which fuse and differentiate into the syncytium during normal tissue turnover [19–23]. In contrast, very little is known at a molecular level about the tegument of cestodes, except for a few individually identified proteins, e.g. [24–27]. Classical studies have shown the presence of transport activities for different nutrients, although these transporters were not identified at a molecular level in most cases [28–30]. Furthermore, methods for obtaining highly enriched fractions of the distal tegument of the cestodes *Echinococcus* spp. and *Hymenolepis* spp. were developed decades ago but their proteins were never identified [31,32]. There is only one recent, low-throughout proteomic analysis of an antigenic fraction enriched in tegumental antigens in *Echinococcus granulosus* [33], using a relatively harsh method for extracting proteins from live worms.

Therefore, it is unclear if there is a common set of tegumental proteins in cestodes, and if these are also shared with other parasitic flatworms. However, members of the “Tegument allergen-like” (TAL) family of proteins, which is specific to parasitic flatworms, have been found in the tegument of different cestodes and trematodes [26,27,34–36]. This family consists of proteins typically containing two N-terminal EF-Hand motifs and a C-terminal dynein light chain (DLC)-like domain. Many different proteins of this family are coded in the genomes of parasitic flatworms, and in some cases, they may lack either conserved domain [27,34]. Although several TAL proteins from trematodes have been characterized structurally and biochemically [37–41], their function is largely unknown.

In this work, we analysed and compared the proteome of the tegument of three cestodes, including larval and adult stages. We identified a common core of tegumental proteins in this group, which also had an important overlap with the described tegumental proteome of trematodes, indicating the existence of an ancestral set of tegumental proteins in neodermatans. Furthermore, we observed differential gene expression among tegumental cytons, evidencing regional specialization within this syncytium.

## Results and Discussion

### Identification of a conserved core of tegumental proteins in cestodes

We have recently adapted to *Mesocestoides corti* the methods described by [31] and [32] for the isolation of the distal tegument of other cestodes, resulting in tegument-enriched fractions (TEFs) [11]. The method is based on mild treatment of live worms with non-ionic detergents at low temperatures, followed by mechanical shearing of the distal tegument by vortexing and isolation by differential centrifugation. In this work, we isolated the distal tegument of larvae (tetrathyridia) of *M. corti*, adults of *Hymenolepis microstoma*, and larvae (protoscoleces) of *Echinococcus multilocularis*. As previously shown for *M. corti* [11], we confirmed by Western Blot in all samples of TEFs a strong enrichment for glycoconjugates that can be detected with the lectin Wheat Germ Agglutinin (WGA), in comparison with whole-worm extracts (WWEs). These glycoconjugates are found in the distal tegument in all three species. In the case of *M. corti*, we also confirmed the enrichment of the tegumental protein alkaline phosphatase (AP) for which we previously developed a specific antibody [11]. In contrast, TEFs were strongly depleted of high molecular weight tropomyosins (HMW-TPMs; [42]), which are markers of muscle fibers, many of which are located directly below the distal tegument. Thus, the isolation procedure results in samples that are highly enriched for the distal tegument (Fig. 1B, C).

Due to the high sensitivity of mass spectrometry methods, one of the main problems for the definition of the proteomes of specific compartments is that some level of contamination with proteins from other compartments is inevitable [43–46]. Therefore, we performed a label-free quantitative proteomic approach, in which we determined the enrichment of each detected protein in tegument-enriched fractions (TEFs) in comparison with whole-worm extracts (WWEs). Conservatively, we defined as tegumental proteins those which were only detected in TEFs in 3 or more replicates, or that had a statistically significant enrichment of 2-fold or higher in TEFs in comparison to WWEs. This stringent approach may therefore exclude some tegumental proteins which are also highly expressed in other tissues.

Following these criteria, we identified 269 tegumental proteins in *M. corti* (including the tegumental marker AP), 287 in *H. microstoma* and 202 in *E. multilocularis* (Tables S1-6). To determine which tegumental proteins are shared among these three cestodes, we first performed a comparison based on BLAST searches, in which we only considered a set of tegumental proteins to be shared among all three species when the proteins were best reciprocal BLASTP hits in all three possible pairwise comparisons between species. In this way, we obtained a core conserved tegumental proteome consisting of 55 proteins shared between the three cestodes (Table 1).

**Table 1.**
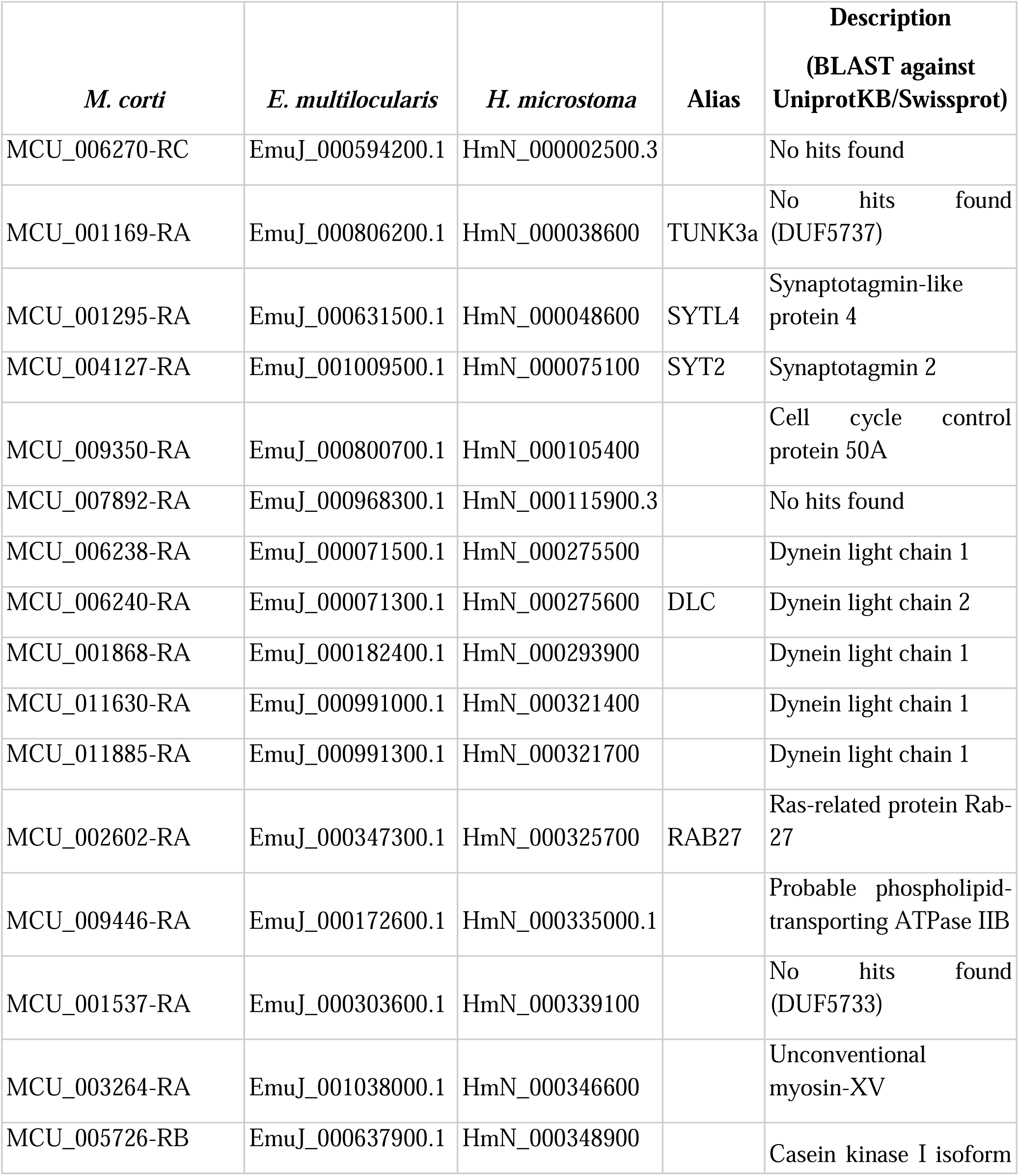
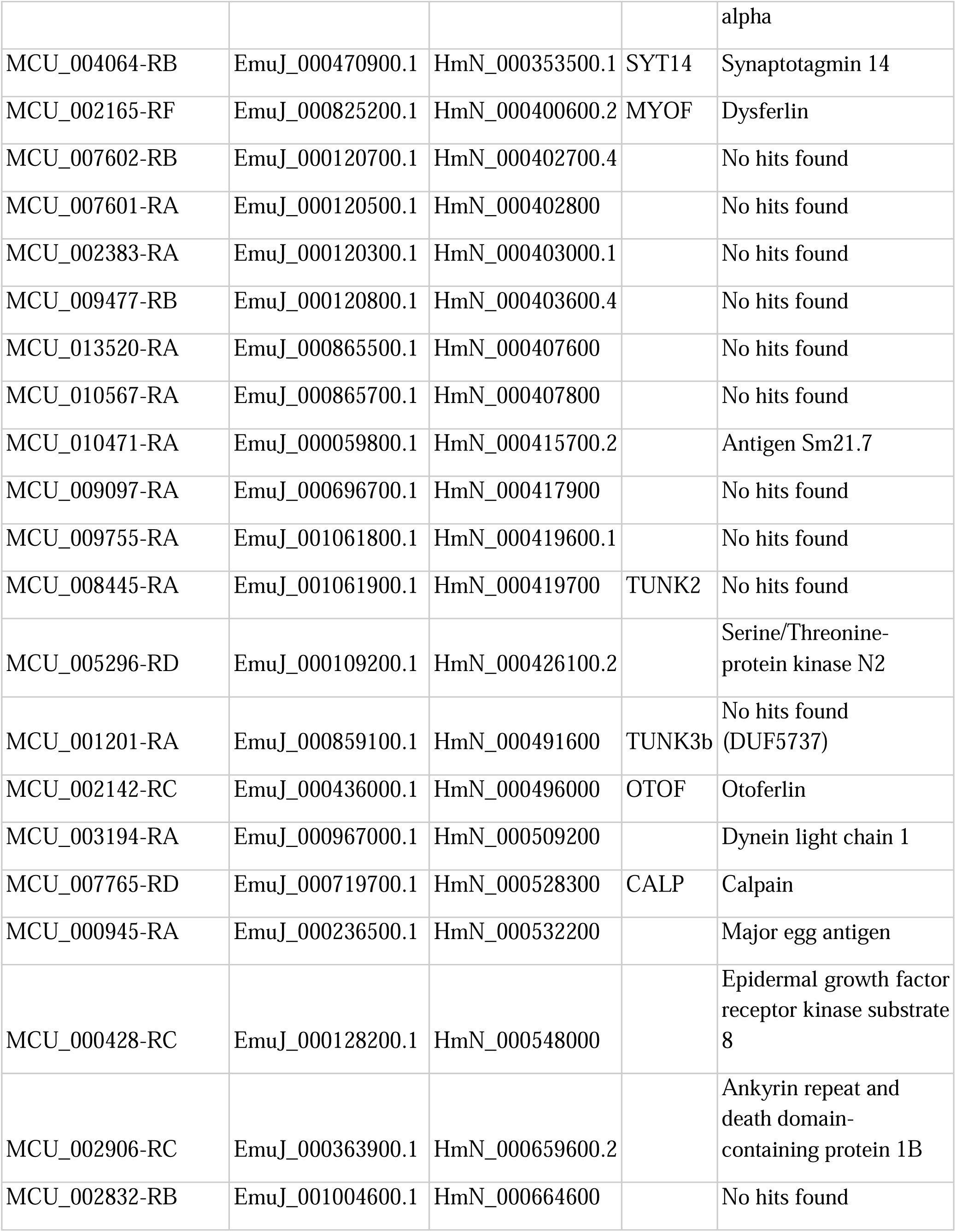
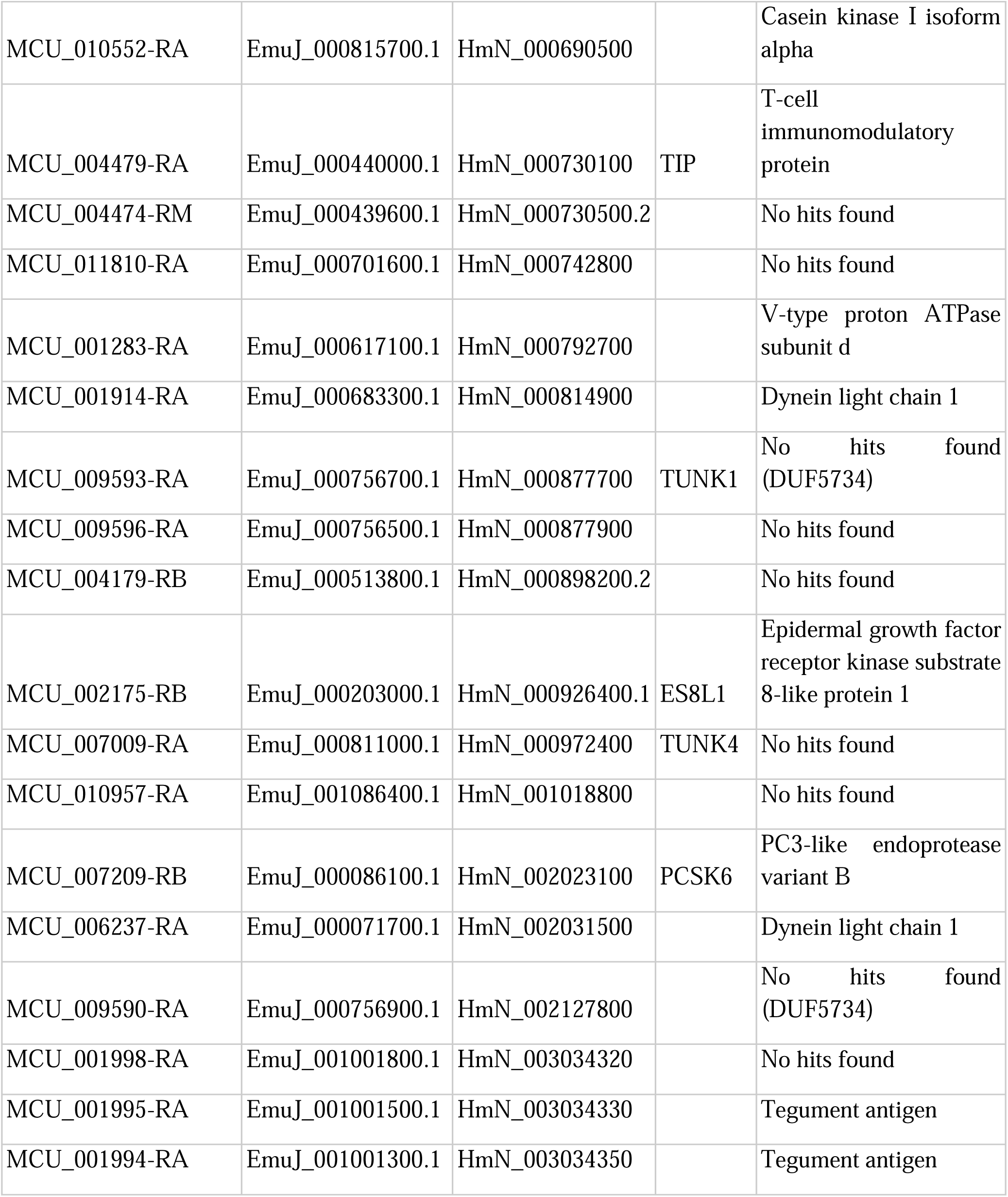
Tegument enriched proteins shared by the three cestode species found by reciprocal BLASTP.

| *M. corti* | *E. multilocularis* | *H. microstoma* | Alias | Description<br>(BLAST against<br>UniprotKB/Swissprot) |
| --- | --- | --- | --- | --- |
| MCU_006270-RC | EmuJ_000594200.1 | HmN_000002500.3 |  | No hits found |
| MCU_001169-RA | EmuJ_000806200.1 | HmN_000038600 | TUNK3a | No hits found<br>(DUF5737) |
| MCU_001295-RA | EmuJ_000631500.1 | HmN_000048600 | SYTL4 | Synaptotagmin-like<br>protein 4 |
| MCU_004127-RA | EmuJ_001009500.1 | HmN_000075100 | SYT2 | Synaptotagmin 2 |
| MCU_009350-RA | EmuJ_000800700.1 | HmN_000105400 |  | Cell cycle control<br>protein 50A |
| MCU_007892-RA | EmuJ_000968300.1 | HmN_000115900.3 |  | No hits found |
| MCU_006238-RA | EmuJ_000071500.1 | HmN_000275500 |  | Dynein light chain 1 |
| MCU_006240-RA | EmuJ_000071300.1 | HmN_000275600 | DLC | Dynein light chain 2 |
| MCU_001868-RA | EmuJ_000182400.1 | HmN_000293900 |  | Dynein light chain 1 |
| MCU_011630-RA | EmuJ_000991000.1 | HmN_000321400 |  | Dynein light chain 1 |
| MCU_011885-RA | EmuJ_000991300.1 | HmN_000321700 |  | Dynein light chain 1 |
| MCU_002602-RA | EmuJ_000347300.1 | HmN_000325700 | RAB27 | Ras-related protein Rab-27 |
| MCU_009446-RA | EmuJ_000172600.1 | HmN_000335000.1 |  | Probable phospholipid-transporting ATPase IIB |
| MCU_001537-RA | EmuJ_000303600.1 | HmN_000339100 |  | No hits found<br>(DUF5733) |
| MCU_003264-RA | EmuJ_001038000.1 | HmN_000346600 |  | Unconventional<br>myosin-XV |
| MCU_005726-RB | EmuJ_000637900.1 | HmN_000348900 |  | Casein kinase I isoform alpha |
| MCU_004064-RB | EmuJ_000470900.1 | HmN_000353500.1 | SYT14 | Synaptotagmin 14 |
| MCU_002165-RF | EmuJ_000825200.1 | HmN_000400600.2 | MYOF | Dysferlin |
| MCU_007602-RB | EmuJ_000120700.1 | HmN_000402700.4 |  | No hits found |
| MCU_007601-RA | EmuJ_000120500.1 | HmN_000402800 |  | No hits found |
| MCU_002383-RA | EmuJ_000120300.1 | HmN_000403000.1 |  | No hits found |
| MCU_009477-RB | EmuJ_000120800.1 | HmN_000403600.4 |  | No hits found |
| MCU_013520-RA | EmuJ_000865500.1 | HmN_000407600 |  | No hits found |
| MCU_010567-RA | EmuJ_000865700.1 | HmN_000407800 |  | No hits found |
| MCU_010471-RA | EmuJ_000059800.1 | HmN_000415700.2 |  | Antigen Sm21.7 |
| MCU_009097-RA | EmuJ_000696700.1 | HmN_000417900 |  | No hits found |
| MCU_009755-RA | EmuJ_001061800.1 | HmN_000419600.1 |  | No hits found |
| MCU_008445-RA | EmuJ_001061900.1 | HmN_000419700 | TUNK2 | No hits found |
| MCU_005296-RD | EmuJ_000109200.1 | HmN_000426100.2 |  | Serine/Threonine-protein kinase N2 |
| MCU_001201-RA | EmuJ_000859100.1 | HmN_000491600 | TUNK3b | No hits found (DUF5737) |
| MCU_002142-RC | EmuJ_000436000.1 | HmN_000496000 | OTOF | Otoferlin |
| MCU_003194-RA | EmuJ_000967000.1 | HmN_000509200 |  | Dynein light chain 1 |
| MCU_007765-RD | EmuJ_000719700.1 | HmN_000528300 | CALP | Calpain |
| MCU_000945-RA | EmuJ_000236500.1 | HmN_000532200 |  | Major egg antigen |
| MCU_000428-RC | EmuJ_000128200.1 | HmN_000548000 |  | Epidermal growth factor receptor kinase substrate 8 |
| MCU_002906-RC | EmuJ_000363900.1 | HmN_000659600.2 |  | Ankyrin repeat and death domain-containing protein 1B |
| MCU_002832-RB | EmuJ_001004600.1 | HmN_000664600 |  | No hits found |
| MCU_010552-RA | EmuJ_000815700.1 | HmN_000690500 |  | Casein kinase I isoform alpha |
| MCU_004479-RA | EmuJ_000440000.1 | HmN_000730100 | TIP | T-cell immunomodulatory protein |
| MCU_004474-RM | EmuJ_000439600.1 | HmN_000730500.2 |  | No hits found |
| MCU_011810-RA | EmuJ_000701600.1 | HmN_000742800 |  | No hits found |
| MCU_001283-RA | EmuJ_000617100.1 | HmN_000792700 |  | V-type proton ATPase subunit d |
| MCU_001914-RA | EmuJ_000683300.1 | HmN_000814900 |  | Dynein light chain 1 |
| MCU_009593-RA | EmuJ_000756700.1 | HmN_000877700 | TUNK1 | No hits found (DUF5734) |
| MCU_009596-RA | EmuJ_000756500.1 | HmN_000877900 |  | No hits found |
| MCU_004179-RB | EmuJ_000513800.1 | HmN_000898200.2 |  | No hits found |
| MCU_002175-RB | EmuJ_000203000.1 | HmN_000926400.1 | ES8L1 | Epidermal growth factor receptor kinase substrate 8-like protein 1 |
| MCU_007009-RA | EmuJ_000811000.1 | HmN_000972400 | TUNK4 | No hits found |
| MCU_010957-RA | EmuJ_001086400.1 | HmN_001018800 |  | No hits found |
| MCU_007209-RB | EmuJ_000086100.1 | HmN_002023100 | PCSK6 | PC3-like endoprotease variant B |
| MCU_006237-RA | EmuJ_000071700.1 | HmN_002031500 |  | Dynein light chain 1 |
| MCU_009590-RA | EmuJ_000756900.1 | HmN_002127800 |  | No hits found (DUF5734) |
| MCU_001998-RA | EmuJ_001001800.1 | HmN_003034320 |  | No hits found |
| MCU_001995-RA | EmuJ_001001500.1 | HmN_003034330 |  | Tegument antigen |
| MCU_001994-RA | EmuJ_001001300.1 | HmN_003034350 |  | Tegument antigen |

As a more stringent approach, we defined Homologs Groups (HGs), [47], with a coverage cutoff of 75% and an identity cutoff of 40%. This resulted in 33 HGs that contained proteins enriched in the tegument of all three cestode species (Table S7), most of which (32/33) were also detected by the reciprocal blast strategy. There was a highly significant overlap of HGs containing tegumental proteins in all pairwise comparison between cestode species (*M. corti* vs *H. microstoma*, 59 shared tegumental HGs among 705 shared detected HGs; p < 0.001; *M. corti* vs *E. multilocularis*, 59 shared tegumental HGs among 779 shared detected HGs; p < 0.001; *H. microstoma* vs *E. multilocularis*, 52 shared tegumental HGs among 1235 shared detected HGs; p < 0.001, Chi-Square test), as well as in the comparison of all three species (33 shared tegumental HGs among 619 shared detected HGs; p < 0.001, Chi-Square test). Altogether, our results show that there is a shared core of tegumental proteins in these species, which was probably present in their last common ancestor. Our proteomic data include both larval and adult stages, indicating that these shared proteins are important for the biology of the tegument in different stages of the life cycle. For *M. corti*, we also compared our results with the proteomic analysis of extracellular vesicles of the same life stage performed by Ancarola et al. [48]. We observed an important overlap in both datasets (30 out of 55 proteins detected by Ancarola et al. were also enriched in our tegumental proteome; Table S8) which indicates the tegumental origin of these extracellular vesicles.

Among the conserved proteins that were enriched in the tegument of the three cestodes species (Table 1), we detected many proteins related to the TAL family, including many short proteins which only have a predicted DLC-like domain. Many conserved tegumental proteins are related to vesicular traffic and the secretory pathway, including homologs of RAB27, Synaptotagmin 2 (SYT2), Synaptotagmin 14 (SYT14), Synaptotagmin like protein 4 (SYTL4), and a member of the Proprotein Convertase family (PCSK6). Other conserved tegumental proteins are related to calcium sensing, including homologs of Myoferlin (MYOF), Otoferlin (OTOF), and a tegument-specific Calpain protease (CALP; see below). Furthermore, several hypothetical proteins without predicted functions are found among the conserved tegumental proteins of cestodes, many of which only have detectable homologs in cestodes or in parasitic flatworms. We have named these as Tegumental Unknown proteins (TUNKs). Some have defined conserved domains of unknown function (DUFs), such as DUF5733, DUF5734 and DUF5737, whereas others have no previously defined domains but have high sequence conservation. In contrast, proteins that were strongly depleted in TEFs in comparison with WWEs include many muscular proteins such as tropomyosin, myosin heavy chain, paramyosin and titin, indicating that contamination with other tissues is low in TEFs (Table S9-14).

In other cases, proteins of the same family were found to be enriched in the tegument of the three cestode species, but members of different HGs were detected in each species, including different annexins, tetraspanins, multidrug resistance transporters, and proteins of the RAB family of GTPases. In order to systematically search for more distant coincidences in the tegumental proteomes of cestodes, we searched for domains that were statistically enriched in the tegument of each species (Table S15). The only domain that was statistically enriched in the tegument of all three species was PF01221 (Dynein light chain type 1), related to the family of TAL proteins. Additionally, other domains are found exclusively or almost exclusively in tegumental proteins in all three species (the low number of proteins containing these domains would prevent them from reaching the threshold for statistical significance). For example, the DUF5734 domain (PF19005) is found almost exclusively in tegumental proteins (five different proteins in *E. multilocularis*, six in *H. microstoma*, and two in *M. corti*), and the genes coding for these proteins are clustered in their genomes.

Finally, our analysis of the proteins enriched in the *H. microstoma* tegument allowed us to identify candidates for well characterized nutrient uptake activities that were described in *Hymenolepis* spp. in previous classic studies [28–30], including a putative sodium-dependent glucose transporter of the SLC5 family (HmN_003033200; SGLT) and a putative sodium dependent concentrative nucleoside transporter (HmN_003032110; CNTA). The ortholog of CNTA was also enriched in the tegument of *M. corti* (MCU_000419). Neither transporter was detected in the tegument of *E. multilocularis* protoscoleces. However, these are quiescent larvae, which were only activated for 3.5 hours before processing them for tegumental isolation. Interestingly, the transcripts coding for these putative transporters in *E. multilocularis* (EmuJ_000714000.1 (SGLT) and EmuJ_000127200 (CNTA)) and *Echinococcus granulosus* (EgrG_000714000 and EgrG_000714050 (SGLT), and EgrG_000127200 (CNTA)) are among the most highly upregulated after protoscolex activation and culture in previous transcriptomic studies [49,50], indicating that these genes and their corresponding transport activities become important once the protoscolex is released from the metacestode vesicles during the infection of the definitive host.

### Some tegumental proteins are released during the tegument isolation procedure

The procedure for the isolation of the distal tegument (which includes a mild detergent treatment and mechanical dissociation) could result in the release of proteins from the distal tegument that are not strongly associated with cellular membranes. This could limit the identification of such tegumental proteins in TEFs. Therefore, we also performed an analysis by quantitative proteomics of the proteins that are released into the supernatants (SNs) during the tegument isolation procedure of *M. corti*, searching for proteins that are enriched in SNs in comparison to the corresponding WWEs (Table S16-19). Many of the proteins that were enriched in TEFs were also enriched in SN samples (95/269 proteins enriched in TEFs were also enriched in SNs, suggesting that these were partially released by the procedure). Other proteins were only enriched in TEFs (174/269 proteins), and these had a higher proportion of proteins with transmembrane domains (11% of proteins have transmembrane domains, compared to only 7% in proteins enriched only in SN), although this difference was not statistically significant (chi-Square Test). Finally, there were 301 proteins which were only enriched in SNs, including several cytosolic and mitochondrial metabolic enzymes, chaperones, and proteins related to the cytoskeleton. Interestingly, of two described fatty acid binding proteins from *M. corti* [51], one was depleted from SNs with respect to WWEs (FABPa; MCU_009121, 2.7-fold depletion) whereas the other was enriched in SNs with respect to WWEs (FABPb; MCU_009116, 3.9-fold enrichment). Neither was enriched in TEFs. Analysis of their localization using specific antibodies showed that both proteins are expressed in many tissues, but only FABPb was present (and enriched) in the distal tegument (Fig. 2). Homologs of known tegumental proteins from other cestodes, including EgGST2 [52] and additional TAL proteins were also found to be enriched in SNs. Therefore, although it is possible that contaminants from other tissues are present, tegumental proteins appear to be enriched in the SN fraction, including some tegumental proteins which could not be identified from TEFs.

**Fig 2.**
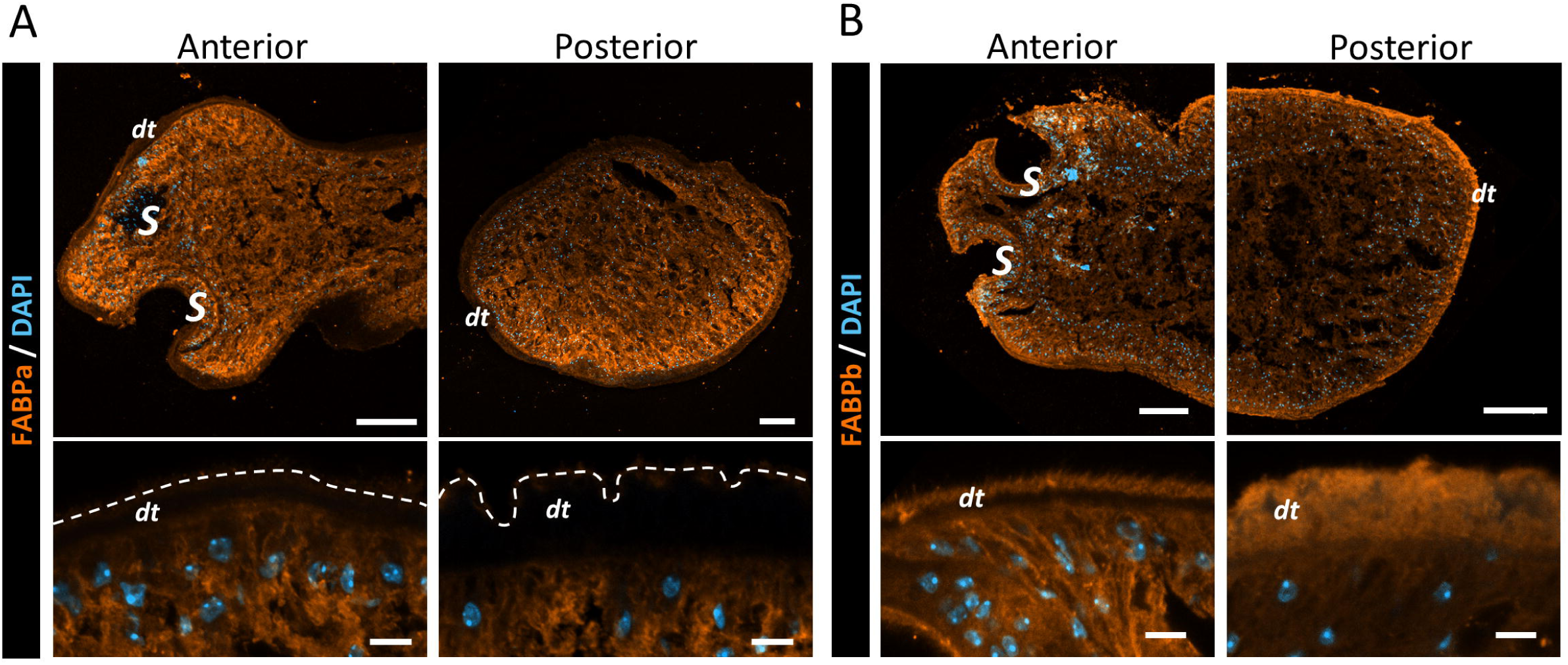
Localization of Fatty Acid Binding Proteins FABPa and FABPb by immunofluorescence in *M. corti*. (A) FABPa is distributed across most *M. corti* tissues, but is absent in the tegument, as shown in detail in the lower panels. (B) FABPb is also present in most tissues, but it is enriched in the tegument, as detailed in the lower panels. Dotted line in the lower panels represents the apical limit of the tegument. dt: distal tegument, S: suckers. Scale bars: upper panels 50 µm, lower panels 5 µm.

### Comparison with the tegumental proteome of the trematode *Schistosoma mansoni*

Next, we compared the tegumental proteins enriched in the tegument of cestodes with the tegumental proteome of the trematode *S. mansoni*, as reviewed by [12] (58 proteins). Homologs of many of these proteins were also described in the tegument of other trematodes [13,14]. This set of proteins was obtained from different studies of isolated tegument samples, which in most cases underwent additional enrichment for integral membrane proteins (for example by differential extraction). Other proteins which were detected in studies of *S. mansoni* but not included in the compilation of Wilson [12] unfortunately could not be included in our comparison, as the original raw data is not available and the proteins were annotated using databases which are no longer extant (reviewed by [8]).

Despite these caveats, we found an overlap between the tegumental proteomes of *S. mansoni* and the three cestodes. Under the more restrictive analysis of HGs, we found four proteins shared between all four species, two of which were also previously confirmed by immunofluorescence to be present in the tegument of *S. mansoni* or *Schistosoma japonicum*: Myoferlin (Smp_141010, [53]), Otoferlin (Smp_163750), Calpain (Smp_214190, formerly Smp_157500, [54]) and a Phospholipid-transporting ATPase (Smp_091650). In the case of Calpain, although more than one paralog from each species is present within this HG, our phylogenetic analysis demonstrates that most tegumental calpains from parasitic flatworms form a single highly supported clade (Fig. S1). Under the more permissive reciprocal blast analysis, other tegumental proteins are found to be shared between *S. mansoni* and the three cestode species, including several TAL proteins (described in greater detail below), RAB27 (Smp_139340) and T-cell immunomodulatory protein (TIP, Smp_194920). Other well characterized tegumental proteins from *S. mansoni* were shared with at least one cestode species, including phosphodiesterase (Smp_153390, [55,56]), tegumental cholinesterase (Smp_136690, [57]), and carbonic anhydrase (Smp_168730, [58]).

Furthermore, other shared tegumental proteins from cestodes have homologs in *S. mansoni* which were not previously detected in the tegument by proteomics, but which are preferentially expressed in tegumental cytons and tegumental progenitors in adult worms according to scRNA-Seq data [19] (in *S. mansoni*, many tegumental proteins are synthesized in the progenitors, and incorporated into the tegument when these fuse with the syncytium [20,59]). These included additional TAL proteins, Synaptotagmin-14 (Smp_150350.1), pro-protein convertase PCSK6 (Smp_149400.1) and a protein with similarity to epidermal growth factor receptor pathway substrate 8 (ES8L1, Smp_139520.1). Finally, based on the available AlphaFold predictions for *E. multilocularis* and *S. mansoni* proteins [60,61], we used FoldSeek [62] to search for structural homologs in *S. mansoni* of cestode TUNKs that did not have detectable similarity by BLAST. Several TUNKs had structural hits in *S. mansoni*, and these in turn showed enriched expression in tegumental cytons or progenitors in the published adult scRNA-Seq dataset, including TUNK1 (DUF5734; best hit Smp_204390, e-6), TUNK2 (no known domains; best hit Smp_121950, e-18), TUNK3 (DUF5737; best hit Smp_075370, e-6) and TUNK4 (no known domains; best hit Smp_126230, e-6).

Altogether, these analyses provide evidence of a core of tegumental proteins that is ancestral to cestodes and trematodes, and therefore likely already present in the last common ancestor of the Neodermata. This core includes several protein families that are specific to parasitic flatworms. It is likely that this shared set of proteins is implicated in basal processes, such as tegumental maintenance and repair. For example, calpains and ferlins, which are found in the tegument of all analyzed parasitic flatworms, have well characterized roles in plasma membrane repair in different eukaryotes [63,64]. This set of shared tegumental proteins between cestodes and *S. mansoni* is relatively small, and it is likely that this is a result of both technical limitations and of higher variability in other tegumental protein families that are directly implicated in host-parasite interactions.

### An expanded complement of tegumental dynein light chains in parasitic flatworms

Several members of the TAL family of proteins have been biochemically characterized from different trematodes [37–41], but very little is known about these proteins in cestodes. For *Echinococcus* spp, two proteins related to TALs were previously shown to be located in the tegument, but each had only either an EF-hand domain [27], or a DLC domain [26]. Classical TAL proteins containing both domains were also found in the genomes of *Echinococcus* spp. [27]. Among the tegument-enriched proteins that we detected in the three species of cestodes, we found many different short proteins containing DLC domains, including divergent proteins for which a DLC domain could be detected but which had no detectable sequence similarity by BLAST (with an e-5 expect value threshold) to known TALs of *Echinococcus* spp. [27].

In order to analyze this set of short and highly divergent proteins which could not be reliably aligned, we performed a similarity clustering analysis using CLANS [65] of all proteins containing DLC domains (Interpro IPR001372) in the genomes of the three cestode species and *S. mansoni* (Fig. 3). Two main groups could be inferred. One group contained the classical TALs, and many of the members of this group also had EF-hand domains. Another group included proteins with high similarity to canonical DLCs of the dynein motor complex, together with many other DLCs with different levels of divergence, none of which had EF-hand domains. Importantly, many proteins from each group were found in the tegumental proteome of each cestode species. Therefore, the complement of tegumental proteins containing DLC domains in parasitic flatworms appears to be much larger and more diverse than previously appreciated.

**Fig 3.**
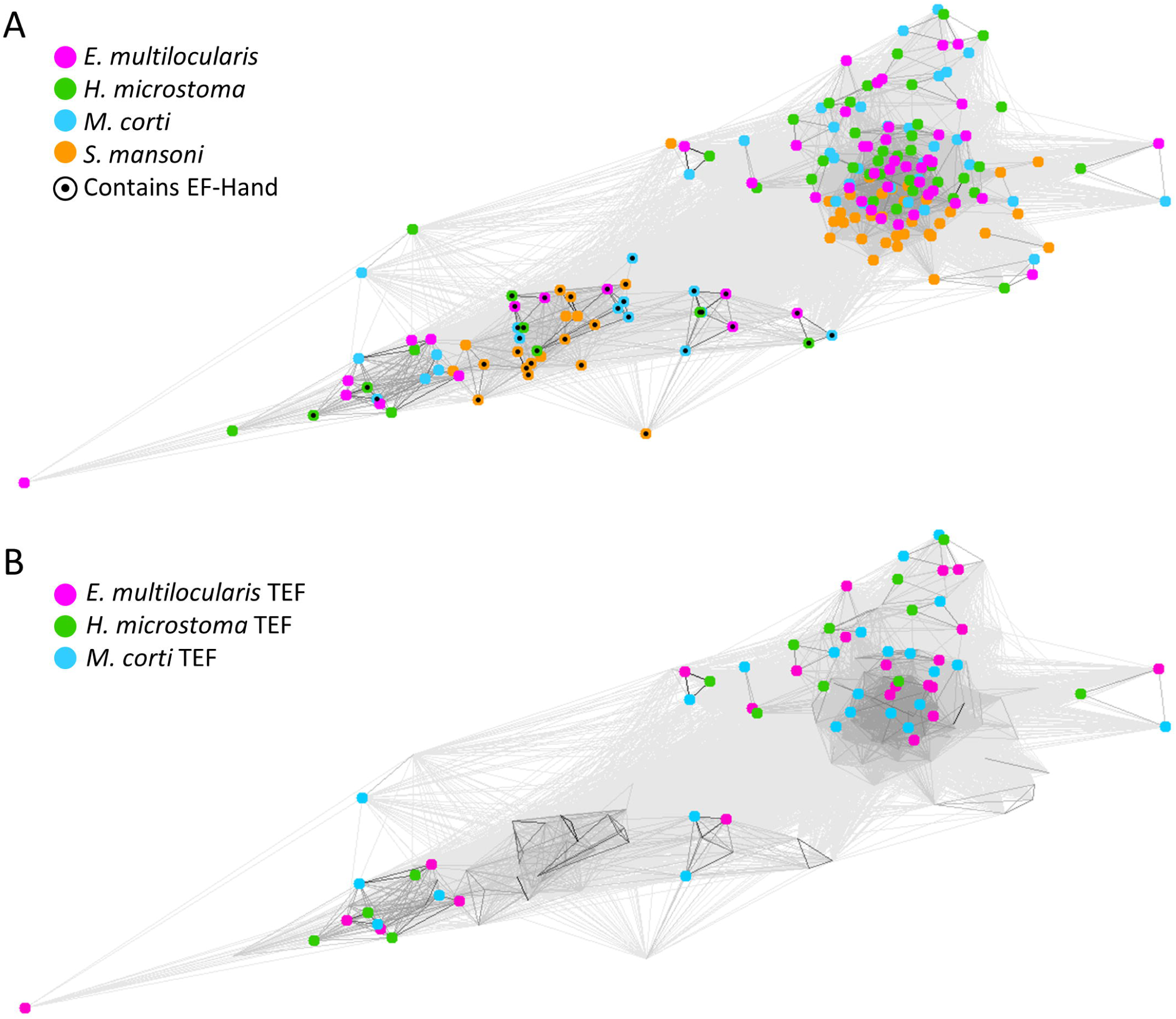
Similarity clustering of proteins containing DLC domains. Each protein is represented by a dot, and lines between dots represent BLASTP hits. (A) Two main groups of DLC containing proteins can be identified in all three cestode species and *S. mansoni*: a tight group (upper right), with no EF-Hand domains, and a second group (lower left), containing the classical TALs, some of which have an EF-Hand domain. (B) Only proteins enriched in TEFs from each species are highlighted. Many proteins from both groups were found to be enriched in TEFs of all three analyzed cestode species.

### Differential expression patterns of tegumental transcripts in cestodes

We analyzed by whole-mount *in situ* hybridization (WMISH) the expression patterns of several genes coding for proteins enriched in the tegument of *H. microstoma* (8 genes) and *E. multilocularis* (6 genes), with sets partially overlapping between both species. We combined this with dextran labeling of the whole tegument [20,11], in order to confirm expression in tegumental cytons (which are interspersed with other cell types, such as muscle and nerve cells, in the sub-tegumental region).

In *E. multilocularis*, all six analyzed genes were expressed in the tegument, and most (5/6) were expressed mainly or exclusively in tegumental cytons (Fig. 4, Table S20). The exception was *em-rab27*, which was widely expressed in most tissues, including in tegumental cytons. Interestingly, *em-calp* was differentially expressed within the tegument, as it was mainly detected in the tegumental cytons of the body but not of the scolex (head). This pattern is reminiscent of the body-specific localization of the tegumental protein emu-TegP11 in protoscoleces [27]. In *H. microstoma*, all eight analyzed genes were expressed mainly or exclusively in the tegument (*hm-syt2* was also detected at lower levels in most tissues) (Fig. 5, S2, Table S20). Expression in the tegument of *hm-myof* is concordant with the recently described expression pattern of its ortholog in *Hymenolepis diminuta* [66] (named as *dysferlin* in that work). Also in this case, we observed differential expression within the tegument, as *hm-calp*, *hm-myof* and *hm-cntA* show little expression in the scolex, and *hm-ap* (coding for an alkaline phosphatase) was completely absent from the scolex except for a crown of tegumental cells in the rostellum (the attachment organ at the anterior end of the scolex) (Fig. 6A, B). These are reminiscent of the rostellar glands of other cestodes, including *Hymenolepis diminuta*, *Hymenolepis nana* and *E. granulosus* which consist of specialized secretory tegumental cytons [67–69]. Furthermore, in the strobila (posterior segmented body) *hm-ap* and *hm-tunk2* are most highly expressed at the posterior end of each segment (Fig. 6A, S2), and expression of *hm-ap* could also be detected in the excretory ducts and in the prostatic glands (Fig. S3). We were unable to obtain antibodies for the protein coded by *hm-ap* that were useful for immunofluorescence, but we could detect alkaline phosphatase activity in the tissues of *H. microstoma* by histochemistry. Remarkably, alkaline phosphatase activity was restricted to the tegument of the strobila and to the rostellum, recapitulating the patterns observed for *hm-ap* by WMISH (Fig. 6C-F). Similarly, alkaline phosphatase activity was highest at the posterior of each segment in the strobila (Fig. 6F), and could also be detected in the excretory ducts (Fig. 6D). Although other alkaline phosphatase genes are present in the genome of *H. microstoma*, our results suggest that differential expression of *hm-ap* between the tegumentary cytons results in the restricted distribution of the enzyme in different regions of the distal tegument syncytium. This is similar to previous results in the related species *H. diminuta*, in which monoclonal antibodies generated towards unknown tegumental proteins showed differential reactivity towards different tegumental cytons and regions of the distal tegument [70]. Similarly, the morphology of the tegument of cestodes has been shown to have regionalized specializations, including different abundance and morphologies of microtriches (microvilli-like projections of the apical membrane) [71–74]. Therefore, although the tegument of cestodes is a giant syncytium, there is clear evidence of regionalized differential gene expression which could be linked to functional and morphological specialization. This is somewhat different to what has been found in adults of the trematode *S. mansoni*, for which differential gene expression in tegumentary cytons is not regionalized, and is thought to result largely from the fusion of different lineages of tegumental progenitors, as they differentiate into mature cytons [59].

**Fig. 4.**
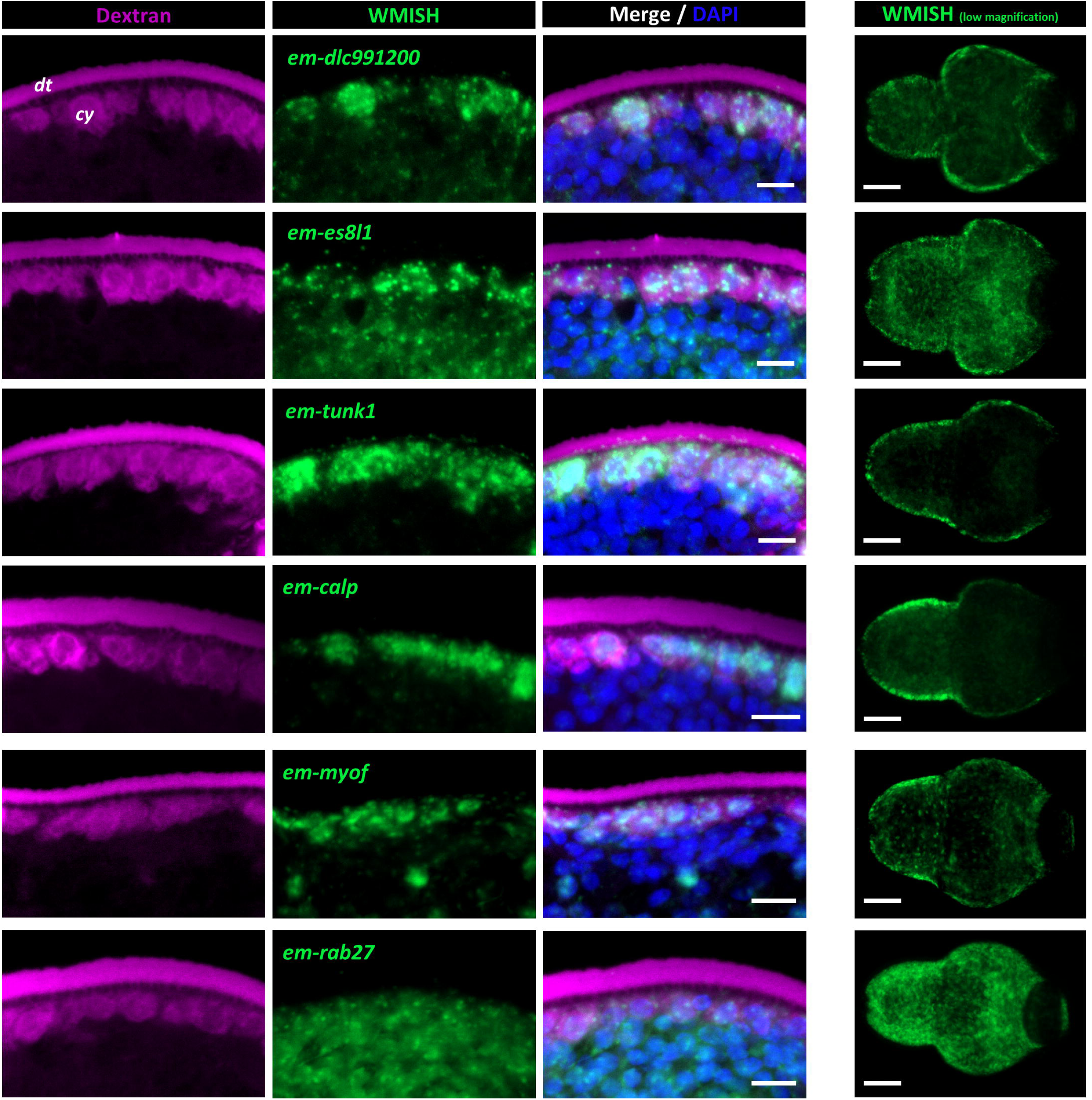
Detection of the expression of tegumental genes by WMISH in *E. multilocularis* protoscoleces. All six genes analyzed colocalize with dextran labeling of the tegument. The right panels show the different global patterns of expression. dt: distal tegument, cy: cytons. Scale bars: merge column: 5 µm, low magnification column: 20 µm.

**Fig. 5.**
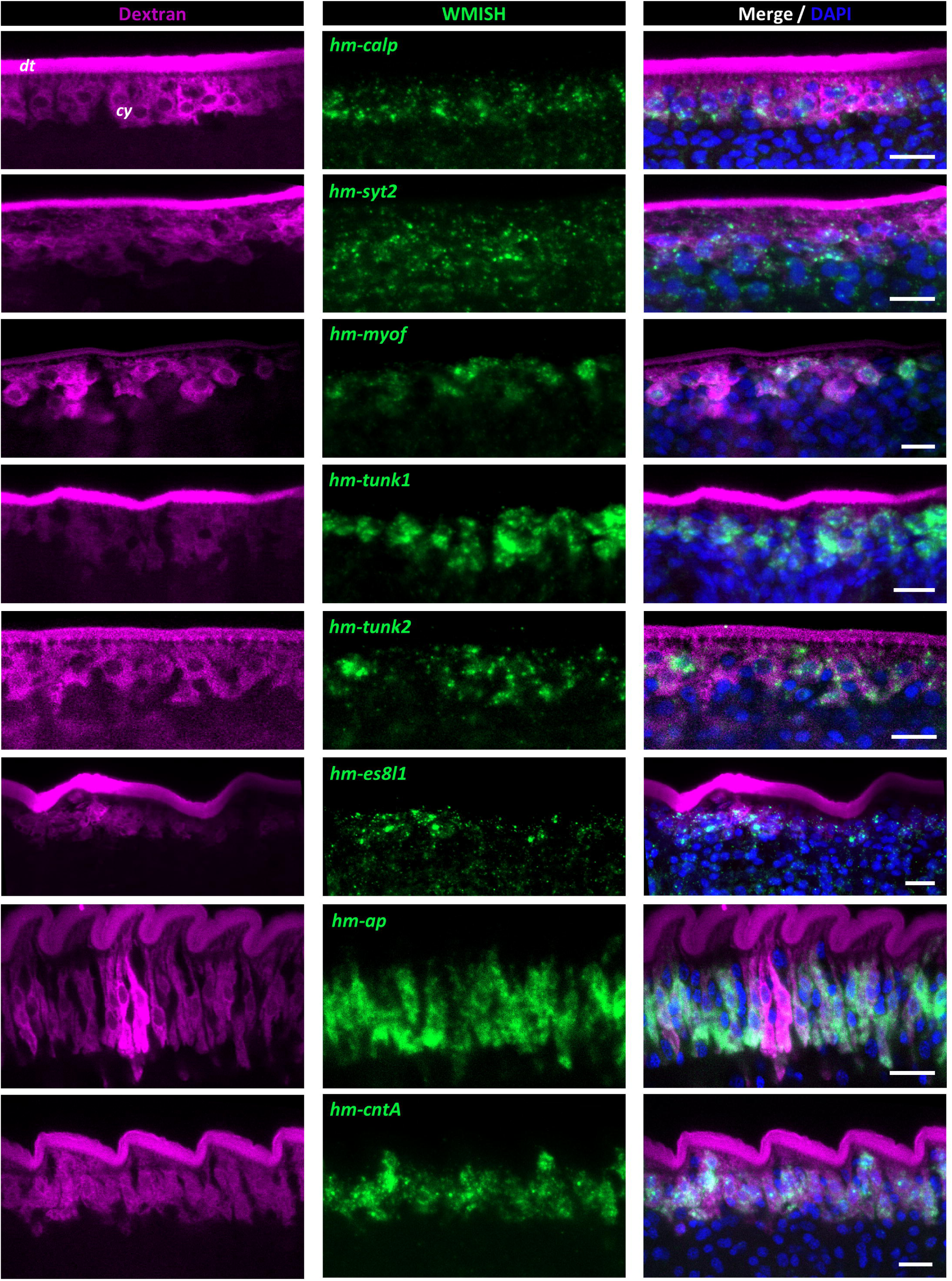
Detection of the expression of tegumental genes by WMISH in *H. microstoma* adults. All eight genes analyzed colocalize with dextran labeling of the tegument. dt: distal tegument, cy: cytons. Scale bars: 10 µm.

**Fig. 6.**
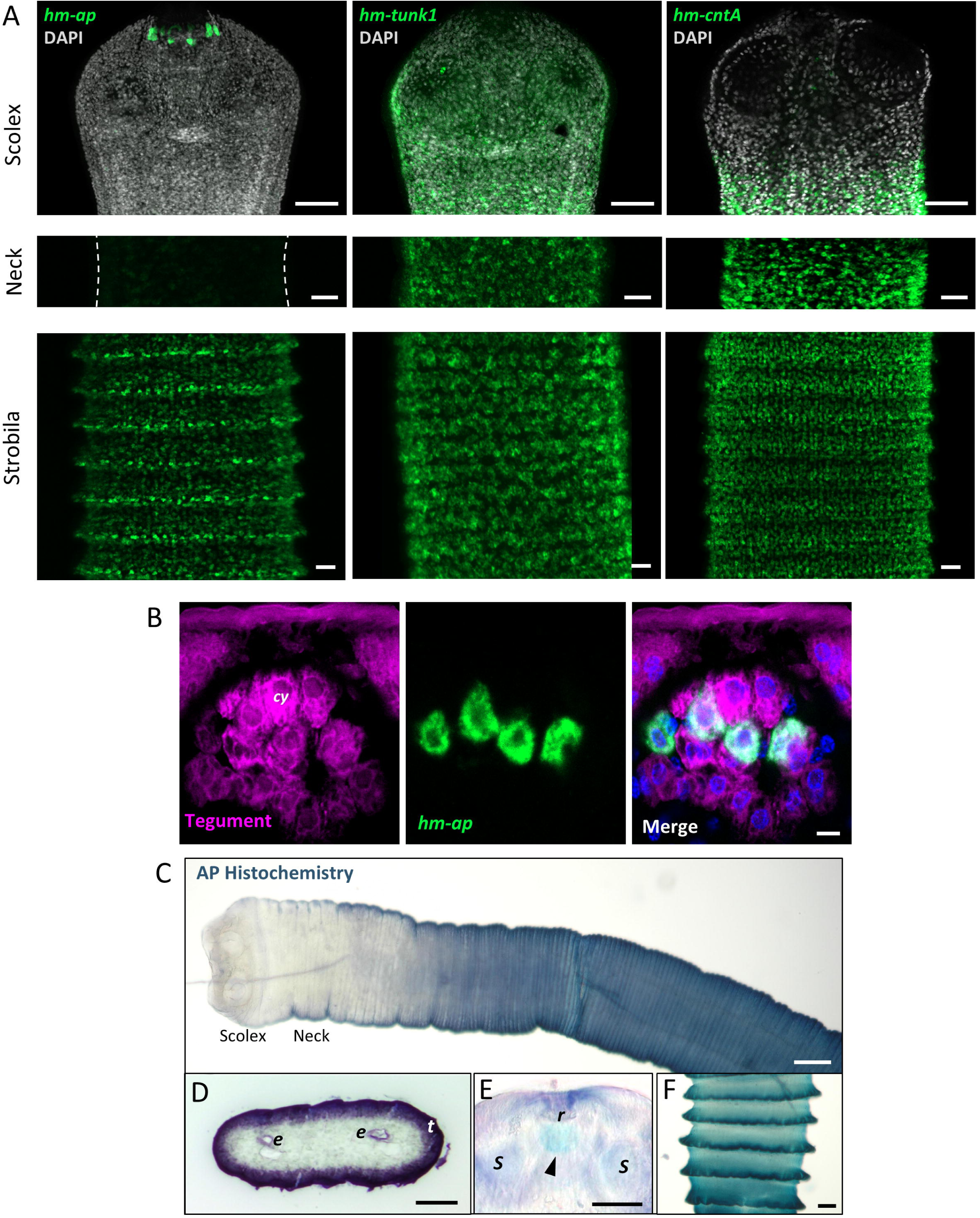
Differential gene expression within the syncytial tegument. (A) Different expression patterns of three tegumental genes: in the scolex, *hm-ap* is only expressed in a few cells surrounding the rostellum, *hm-tunk1* is expressed widely, and *hm-cntA* is completely absent. In the neck, *hm-tunk1* and *hm-cntA* are expressed strongly, whereas *hm-ap* is barely detected in this region. In the strobila, expression of *hm-ap* increases progressively towards the posterior regions, where the signal is stronger in the posterior edge of each segment, whereas *hm-tunk1* and *hm-cntA* show uniform expression in the tegument in this region. (B) Cells expressing *hm-ap* surrounding the rostellum are connected to the tegument as shown by colocalization with dextran labeling (tegument). (C) Detection of alkaline phosphatase (AP) activity by histochemistry is correlated to *hm-ap* expression. (D) Histochemistry on a transversal section of the strobila showing AP activity in the tegument and excretory ducts. (E) Signal surrounding the rostellum was also detected by AP histochemistry. (F) AP activity is higher in the posterior region of each segment cy: cytons, dt: distal tegument, e: excretory ducts, t: tegument, r: rostellum, s: suckers. Scale bars: A: scolex 50 µm, middle and lower panels 20 µm; B: 5 µm, C-F: 50 µm.

In conclusion, this work provides the first comprehensive proteomic characterization of the tegument of cestodes, comparing multiple species and life stages. We demonstrate that many tegumental components are shared not only among cestodes but also with the trematode *Schistosoma mansoni*, supporting the existence of an ancestral tegumental proteome, including conserved tegumental proteins that are likely specific to parasitic flatworms. Furthermore, the demonstration of regionally restricted gene expression within tegumental cytons indicates functional specialization within this syncytium. Together, these findings advance our understanding of tegument biology and evolution, and establish a framework for future studies towards the identification of new targets against cestode infections.

## Materials and Methods

### Parasite material

*M. corti* (syn. *M. vogae*) tetrathyridia larvae from the strain originally isolated by Specht and Vogue [75] were obtained from intraperitoneal infected C57BL/6 mice, in collaboration with the Laboratorio de Experimentación Animal, Facultad de Química, Universidad de la República, Uruguay (“Mantenimiento del cestodo *Mesocestoides vogae* para estudios de actividad antihelmíntica y uso de otros servicios de Udelar”; protocol number 10190000011119, approved by Comisión Honoraria de Experimentación Animal, Uruguay). Mice were euthanized by cervical dislocation, and the parasites were collected under sterile conditions at three to six months post-infection.

The life cycle of *H. microstoma* was maintained using C57BL/6 mice as definitive hosts, and the beetle *Tribolium confusum* as intermediate hosts [76], also in collaboration with the Laboratorio de Experimentación Animal, Facultad de Química, Universidad de la República, Uruguay (“Mantenimiento del ciclo vital completo del cestodo *H. microstoma* utilizando sus hospedadores naturales *Mus musculus* (ratón) y *Tribolium confusum* (escarabajo de la harina)”, protocol number 10190000025215, approved by Comisión Honoraria de Experimentación Animal, Uruguay). Infected mice were euthanized by cervical dislocation, adult worms were obtained from the bile duct and intestine and kept in Phosphated Buffered Saline (PBS) briefly until further use for tegument isolation experiments or for fixation.

Experiments with *E. multilocularis* were carried out using parasite isolates that derive from Old World Monkey species naturally infected in a breeding enclosure [77]. Metacestode tissue was propagated by intraperitoneal passage in Mongolian jirds (*Meriones unguiculatus*) as previously described [78]. Protoscoleces were isolated from metacestode tissues and activated by low pH/pepsin/taurocholate treatment as described by Ritler et al. [79]. The animal study was approved by the Ethics Committee of the Government of Lower Franconia, Würzburg, Germany, under permit numbers 55.2–2531.01-61/13 and 55.2.2-2532-2-1479-8.

### *In vitro* culture of *M. corti* tetrathyridia

After collection, larvae were kept in PBS at 4°C until use (up to 7 days). Before conducting the isolation of tegumental fractions, larvae were cultured for 12-24 h in RPMI-1640 medium (Sigma, R4130, United Kingdom) supplemented with 10% Fetal Bovine Serum (Capricorn, cat N° FBS-11A, collected in South America) and antibiotic-antimycotic (Thermo, 15240–062, U.S.A.), at 37°C and 5% CO2.

### Tegument isolation and total protein extracts

In order to isolate the tegument and obtain tegument enriched fractions, we followed our previously published protocol [11], adapted from [31,32]. Briefly, one volume of parasites was transferred to three volumes of PBS + 0.2% Triton-X 100 (Baker, U.S.A.), put on ice and mild shaking for 10 minutes (Thermolyne, Roto Mix type 50800, at minimal speed). Afterwards, tubes containing the parasites were vortexed for 30 seconds in order to mechanically detach the distal tegument. Differential centrifugation was conducted at 4°C, first at 200 g for 3 minutes, obtaining a pellet composed by the “naked” worms. The supernatant was recovered and centrifuged at 2500 g at 4°C for 15 minutes. After this second centrifugation, the supernatant was removed and PBS was added to wash the pellet, followed by another centrifugation at 2500 g for 15 minutes. The supernatant from the washing step was discarded, removing as much liquid as possible, and the resulting pellets, corresponding to the tegument-enriched fractions (TEFs), were frozen at -80°C until further use. TEF pellets were dissolved in an incomplete Loading Buffer: 75 mM Tris pH 6.8, 0.6% SDS, 15% glycerol (without β-mercaptoethanol and Bromophenol blue) plus protease inhibitor cocktail (Sigma, P8849, Germany) and boiled at 100°C for 10 minutes.

For total protein extracts from whole worms (WWEs), parasites were quickly frozen in liquid nitrogen. Afterwards, two volumes of incomplete Loading Buffer (without β-mercaptoethanol and Bromophenol blue) plus protease inhibitor cocktail (Sigma, P8849, Germany) were added and parasites were put on ice and disrupted with a pestle homogenizer for 5 minutes. The resulting homogenate was boiled at 100°C for 10 minutes and then centrifugated at room temperature at 17.000 g for 5 minutes. The supernatant was carefully recovered and stored at - 80°C until further use.

Protein quantification of TEFs and WWEs was conducted using the Pierce BCA Protein Assay Kit (Thermo, 23227, U.S.A.). After quantification, β-mercaptoethanol and Bromophenol blue were added to the extracts to final concentrations of 7.5% and 10 μg/mL, boiled at 100°C for 10 minutes and stored at -80 °C until further use. For the analysis of proteins released into the supernatant in *M. corti* (SN), the supernatant obtained during tegument isolation after centrifugation at 2500 g was collected and centrifuged at 20.000 g at 4 °C for 15 minutes. The supernatant obtained in this last centrifugation step was quantified as described above, and stored at -80 °C until further use.

### Western Blot

Samples (20 μg) were run in a 10% SDS-PAGE gel. Proteins were then transferred to a nitrocellulose membrane (Thermo, 88018, Germany). The membrane was blocked with 5% milk prepared in Tris-NaCl plus 0.1% Tween-20 (Sigma, P9416, U.S.A.) (TBS-T). After blocking, the membrane was incubated overnight with anti-high molecular weight tropomyosin antibody [42] or anti-*M. corti* AP antibody [11] in blocking solution at 4°C with rocking. After washing with TBS-T, the membrane was incubated with a secondary antibody conjugated to Horseradish Peroxidase (Thermo, A16104) in blocking solution at room temperature for 1 h with rocking. Finally, the membrane was washed with TBS-T and developed using the chemiluminescent reagent SuperSignal West Pico PLUS (Thermo, 34580, U.S.A.). Imaging was performed with a G:BOX system.

For the detection of glycoconjugates, the membrane was incubated with lectin WGA conjugated to rhodamine (Vector Labs, U.S.A.) in TBS-T for 1 h at room temperature with rocking, in the dark. Imaging was done with a G:BOX system.

### Protein sample preparation for Mass Spectrometry analysis

Proteomic analysis of WWEs and TEFs was carried out using four biological replicates (for *H. microstoma* and *M. corti*) or three replicates (for *E. multilocularis*). For each sample, 20 μg of protein were loaded onto a 12.5% acrylamide gel and subjected to electrophoresis until proteins migrated approximately 1 cm into the resolving gel. Gels were subsequently fixed and stained with Coomassie Blue. Gel fragments were manually excised and transferred to microcentrifuge tubes.

Sample processing for mass spectrometry analysis included protein reduction with 10 mM dithiothreitol at 56 °C for 1 h under agitation; cysteine alkylation with 50 mM iodoacetamide at room temperature for 45 min in the dark with agitation; *in-gel* protein digestion was carried out overnight at 37 °C using 0.1 mg/mL sequencing-grade trypsin (Promega) in 50 mM ammonium bicarbonate; peptides were extracted by incubation with 60% acetonitrile/0.1% trifluoroacetic acid for 2 h at 30 °C with vigorous agitation; and finally, samples were dried under vacuum and resuspended in 0.1% trifluoroacetic acid.

Peptide samples were desalted using C18 micro-columns (OMIX C18 pipette tips, Agilent), followed by elution with 0.1% formic acid in acetonitrile, vacuum drying, and resuspension in 0.1% formic acid. Peptide concentration was measured at 215 nm using a Denovix DS-11 FX+ spectrophotometer/fluorometer. The final volume was adjusted to normalize sample concentrations.

### Nanoscale liquid chromatography-tandem mass spectrometry analysis

A shotgun proteomic strategy was used to compare protein profiles of WWEs and TEFs within each studied organism. Samples from *M. corti* and *H. microstoma* were analyzed using a Q Exactive Plus mass spectrometer, whereas *E. multilocularis* samples were analyzed using an Orbitrap Exploris 240. In all cases, a nano-HPLC Ultimate 3000 system (Thermo Scientific) was coupled online to the mass spectrometer via an Easy-Spray source.

Five μg of tryptic peptides were injected into an Acclaim PepMap^TM^100 C18 nano-trap column (75 μm x 2 cm, 3 μm particle size), and separated using a 2 μm particle size, 75 mm x 500 mm, Easy-Spray^TM^ analytical C18 HPLC column at a constant flow rate of 200 nL/min and 40 °C and applying the following gradient: from 1% to 35% mobile phase B (0.1% formic acid in acetonitrile) over 150 min, followed by 35% to 99% mobile phase B over 20 min.

The mass spectrometer was operated in positive ion mode with a spray voltage of 2.0 kV and a capillary temperature of 250 °C. Data-dependent acquisition was performed in two stages. Full MS scans were acquired over an m/z range of 200–2000, with a resolution of 90,000 (Orbitrap Exploris 240) or 70,000 (Q Exactive Plus). The top 12 most intense precursor ions were selected for HCD fragmentation using stepped normalized collision energies of 25, 30, and 35, with MS/MS resolution set to 25,000 (Exploris 240) or 17,500 (Q Exactive Plus). Precursor ions with charge states between 2 and 5 (Exploris 240) or 2 and 4 (Q Exactive Plus) were selected for fragmentation. Dynamic exclusion was set to 10 s (Exploris 240) or 5 s (Q Exactive Plus).

### Proteomic data analysis

Database generation, protein identification, and downstream analyses were performed using PatternLab V (https://www.patternlabforproteomics.org) [80]. For this purpose, protein sequence datasets for *M. corti*, *H. microstoma*, and *E. multilocularis* were retrieved from WormBase ParaSite (https://parasite.wormbase.org/) in May 2022, August 2022, and April 2024, respectively. Because the predicted proteome of *M. corti* had a very large number of isoforms with little support for many genes, we pruned this database to a single isoform per gene, according to the following hierarchical criteria. First, we searched for evidence of each isoform in an independent preliminary proteomic dataset of TEFs and WWEs (two samples from each), selecting the isoform that was present in the largest number of samples, and in case of a tie, further selecting the isoform with the highest protein score. Second, if no isoform was detected in the preliminary proteomic dataset, we selected the isoform with the highest BLASTP bitscore against the UniProtKB database. Third, if no BLASTP hit was detected, we selected the longest isoform. In addition, sequences corresponding to the 127 most common mass spectrometry contaminants were included to construct a target-decoy database for each organism.

Raw files were searched against the database using the Comet search engine (integrated in PatternLab V), applying the following parameters: fully specific trypsin as the proteolytic enzyme, allowing up to two missed cleavages; methionine oxidation as a variable modification and cysteine carbamidomethylation as a fixed modification; and a precursor mass tolerance of 35 ppm. Peptide-spectrum matches were filtered using the Search Engine Processor, applying stringent FDR criteria set at 3% at the spectrum level, 2% at the peptide level, and 1% at the protein level. To identify proteins exclusively detected in each sample set (WWEs and TEFs) within each organism, PatternLab’s approximately area-proportional Venn diagram module was used with a significance threshold of p ≤ 0.05. To detect proteins present in both conditions but showing significantly different relative abundances, the pairwise comparison module was applied based on extracted ion chromatogram (XIC) analysis, using the following statistical criteria: |log fold change| ≥ 1 and p ≤ 0.05.

### Data availability

The mass spectrometry proteomic raw data have been deposited to the ProteomeXchange Consortium via the PRIDE partner repository with the dataset identifier PXD077018 (doi: 10.1093/nar/gkae1011.)

### Comparative analysis of tegumental proteomes

Putative orthologs among the proteins enriched in the tegument of *H. microstoma*, *M. corti* and *E. multilocularis* were identified by a three-way reciprocal BLASTP hit strategy [81]. Proteins were considered orthologs only if they were best reciprocal BLASTP hits in all three possible pairwise comparisons between species.

For the generation of more stringent Homologs Groups (HGs), the predicted proteomes from six cestodes species (*Echinococcus canadensis, Echinococcus granulosus, Echinococcus multilocularis, Hymenolepis microstoma* and *Hymenolepis nana*, and *M. corti*, pruned to a single isoform per gene as described above) and two trematode species (*Clonorchis sinensis* and *Schistosoma mansoni*) were downloaded from WormBase Parasite (2023/04/28) [82]. These were used to construct HGs using the software get_homologues [47] with default parameters except for: Bidirectional Best Hit algorithm (BDBH) and minimal identity percentage of 40%, executed locally.

Bash scripting was used to identify which HGs contained proteins detected by mass spectrometry for each of the species and samples analysed, and to evaluate the overlap between the three cestode species, as well as the overlap with the *S. mansoni* tegumental proteome [12]. Significance in the overlap of tegument enriched HGs of cestodes was assessed using Chi-Square test performed on Past4.0 software [83].

In order to obtain a list of the protein domains present in our proteomic data, batch CD-Search against the Pfam database [84] was conducted and enriched domains were identified performing Fisher’s test with Benjamini-Hochberg correction in R [85].

### Structural search of distant homologs in *S. mansoni*

In order to identify distant *S. mansoni* homologs of cestode TUNKs, we performed a structural homology search. AlphaFold predictions of *E. multilocularis* proteins were downloaded in PDB format from the AlphaFold Database (alphafold.ebi.ac.uk) [60,61], and used to search for similar structures in the AlphaFold Database with FoldSeek (search.foldseek.com/) [62]. In those cases where a *S. mansoni* hit was obtained, the expression pattern of the corresponding gene was retrieved from the scRNA-Seq dataset of adult *S. mansoni* [19].

### Phylogenetic analysis of calpains

Calpain sequences of HG 28044_egra.2211.faa were aligned with MUSCLE [86] and Maximum Likelihood Phylogenetic inference was performed with MEGA11 [87]. The evolutionary model LG + G + I was selected based on the Find Best Protein Model feature.

### Analysis of proteins containing Dynein Light Chain domains

Proteins containing the DLC domain (InterPro IPR001372) in *E. multilocularis*, *H. microstoma*, *M. corti* and *S. mansoni* were downloaded from WormBase Parasite [82] via the Biomart tool. EF-Hand domains (InterPro IPR002048 and IPR011992) from these proteins were also extracted from their WormBase Parasite annotations. Clustering was performed using CLANS [65], which is based on pairwise sequence similarity from BLASTP comparisons, with an e-value cutoff of e-5.

### Fluorescent labeling of the tegument with Dextran-Rhodamine

The tegument was fluorescently labeled with dextran-rhodamine (Thermo, D3312, U.S.A.) as described by Guarnaschelli and Koziol [11] for *M. corti*, based on the protocol originally described for *S. mansoni* by Wendt et al [20]. Briefly, *H. microstoma* adults and *E. multilocularis* protoscolices were incubated in 5 volumes of a solution of distilled water with 2 mg/mL dextran in a tube wrapped in aluminum foil, and vortexed for 3 minutes at 50% intensity. The dextran solution was immediately removed and parasites were quickly washed with PBS three times, after which they were fixed with 4% paraformaldehyde in PBS overnight. Parasites were then transferred to methanol and stored at -20°C until further use.

### Whole Mount *in situ* Hybridization

Fragments of cDNA of each gene of interest were amplified using the primers listed in Table S21. Amplicons were cloned into pCR II vectors (Dual promoter Kit, Invitrogen, 45–0007LT, U.S.A.) and digoxigenin-labeled RNA probes were synthesized by *in vitro* transcription using SP6 or T7 polymerases (Thermo, EP0131 and EP0111 respectively), using 3.5 mM digoxigenin-UTP, 6.5 mM UTP, and 10 mM CTP, GTP and ATP. Whole mount *in situ* hybridization was conducted as described by Koziol et al [88], in samples of paraformaldehyde-fixed worms with and without dextran labelling of the tegument.

Experiments were imaged by confocal microscopy (Zeiss LSM 800CyAn, Advanced Bioimaging Unit of the Institut Pasteur of Montevideo).

### Immunofluorescence

Experiments were conducted on cryosections of *M. corti* tetrathyridia larvae and *H. microstoma* adults, and on paraplast sections of *E. multilocularis* protoscoleces adapting the protocols published by Koziol et al [89]. Cryosections and rehydrated paraplast sections were washed with PBS with 0.1% Triton X-100 (Baker, USA) (PBS-T) and then blocked with PBS-T plus 1% Bovine Serum Albumin (BSA; Capricorn, USA) and 5% sheep serum (Sigma, S3772, USA). Incubations with the primary and secondary antibodies were done for 72 hours each, in PBS-T with 1% BSA and 0.02% sodium azide. This extended incubation was necessary to ensure the adequate penetration of the antibodies in the distal tegument. After incubation with each antibody, slides were extensively washed with PBS-T. Staining with 4’,6-diamidino-phenylindole (DAPI) was performed together with the secondary antibody incubation. The specimens were mounted with Fluoroshield (Sigma, F6182, U.S.A.). The purified antisera used to detect FABPa and FABPb were kindly provided by Gabriela Alvite [51], and used in a 1 in 40 dilution; the rabbit polyclonal antibody against high molecular weight tropomyosin isoforms [42] was used in a 1 in 500 dilution. Anti-rabbit secondary antibody conjugated to Alexa 546 (Thermo, A11010, U.S.A.) was used in a 1 in 1000 dilution. For detection of glycoconjugates, sections were washed with PBS and incubated in the dark for 72 hours with PBS-T plus 1% BSA and 10 μg/mL fluorescent Wheat Germ Agglutinin (WGA, Vector Labs, USA), DAPI and 0.02% sodium azide. The slides were washed extensively with PBS-T and mounted with Fluoroshield.

### Alkaline phosphatase histochemistry

Alkaline phosphatase histochemistry was performed in cryosections and whole-mounts of *H. microstoma* adults with nitro blue tetrazolium chloride and 5-bromo-4-chloro-3-indolyl phosphate as previously described [90].

## Acknowledgements

The authors thank Beatriz Munguía, Laboratorio de Experimentación Animal, Facultad de Química, Universidad de la República, Uruguay for providing *M. corti* larvae and maintaining mice infected with *H. microstoma* adults; and Gabriela Alvite, Sección Bioquímica, Facultad de Ciencias, Universidad de la República, for kindly providing the anti-FABPa and anti-FABPb antibodies. The authors gratefully acknowledge the Advanced Bioimaging Unit and the Unidad de Bioquímica y Proteómica Analíticas at the Institut Pasteur Montevideo for their support and assistance in the present work. This work was funded by Agencia Nacional de Investigación e Innovación (ANII), Uruguay (project FCE_1_2021_1_166701 to UK and POS_FCE_2021_1_1010808 PhD Fellowship to IG), Comisión Académica de Posgrados, Universidad de la República, Uruguay (PhD Fellowship to IG) and Programa de Desarrollo de las Ciencias Básicas (PEDECIBA), Uruguay. AI, AL, MP, JC, and UK are members of Sistema Nacional de Investigadores (SNI-ANII).

## Author contributions

IG, AL, KB, MCR, AI, UK planned the experiments. IG, AL, RV, MB, MP, MC, UK performed the experiments. IG, AL, JC, AI, UK analysed the data. KB contributed reagents or other essential material. IG, UK wrote the paper.

## Data availability statement

The mass spectrometry proteomic raw data have been deposited to the ProteomeXchange Consortium via the PRIDE partner repository with the dataset identifier PXD077018 (ref: doi: 10.1093/nar/gkae1011.). Additional data that support the findings of this study are available in the article and its Supporting Information. Additional raw data is available under request from the corresponding authors.

## Abbreviations

AP: alkaline phosphatase
BSA: Bovine Serum Albumin
DAPI: 4’,6-diamidino-phenylindole
DLC: dynein light chain
DUF: domain of unknown function
HG: Homolog Group
HMW-TPM: High molecular weight tropomyosin
PBS: Phosphated Buffered Saline
scRNA-Seq: single cell RNA sequencing
SN: supernatant
TAL: Tegument Allergen-Like
TBS-T: Tris-Buffered Saline plus 0.1% Tween-20
TEF: tegument-enriched fraction
TUNK: tegumental unknown protein
WGA: Wheat Germ Agglutinin
WMISH: whole-mount in situ hybridization
WWE: whole-worm extract

