## Supplementary Figure S1 for "Comparative proteomics reveals a conserved core of tegumental proteins in parasitic flatworms"

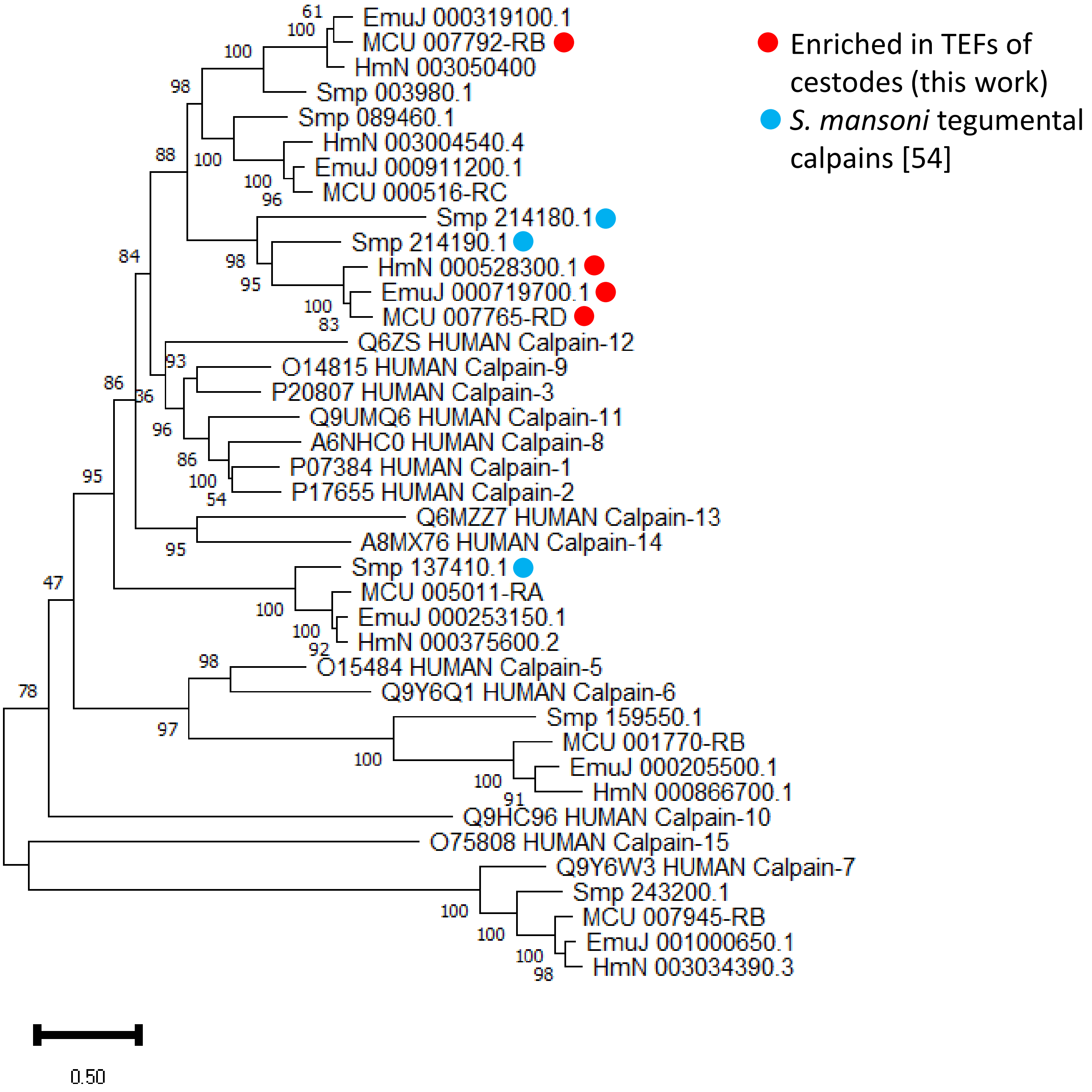


Fig. S1. Phylogeny of calpains. Calpains from *E. multilocularis*, *H. microstoma*, *M. corti*, *S. mansoni* and *Homo sapiens* were used to construct a phylogenetic tree using the Maximum Likelihood method (LG + G + I model). Tegumental calpains form a well supported clade
