## Supplementary Figure S2 for "Comparative proteomics reveals a conserved core of tegumental proteins in parasitic flatworms"

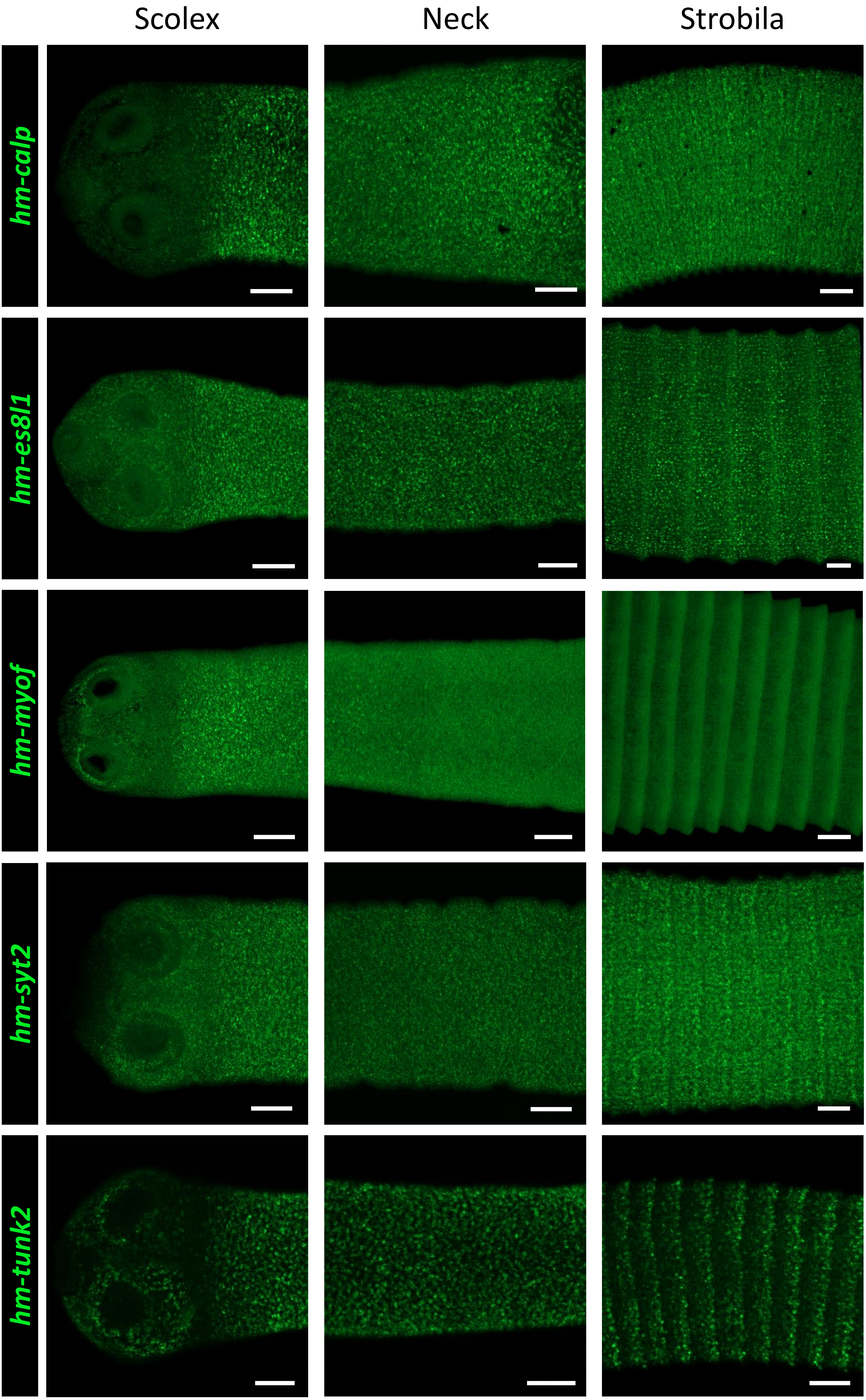

Scolex
Neck
Strobila
hm-calp
hm-es8l1
hm-myof
hm-syt2
hm-tunk2

Fig. S2. Differential expression of additional genes within the syncytial tegument. WMISH shows different spatial distribution of gene expression among tegumental genes. *hm-calp* and *hm-myof* expression is very low or absent in most of the scolex. *hm-es8l1*, *hm-syt2* and *hm-tunk2* are strongly expressed in scolex, neck and strobila. Scale bars: 50 µm
