## Supplementary Figure S3 for "Comparative proteomics reveals a conserved core of tegumental proteins in parasitic flatworms"

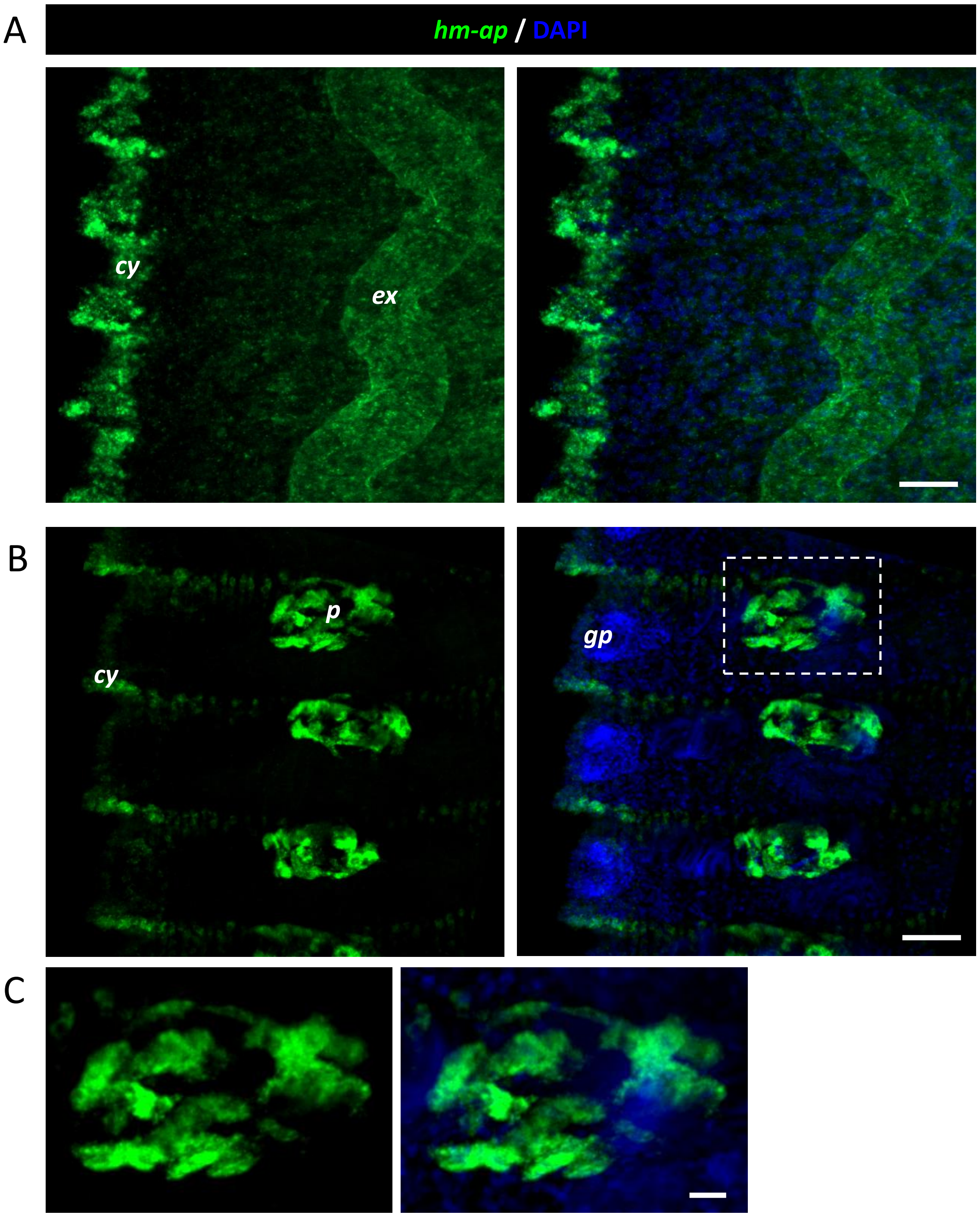


Fig. S3. Additional domains of expression of *hm-ap* detected by whole mount *in situ* hybridization. In addition to the tegument, *hm-ap* was also detected in the excretory ducts (A) and in the prostatic glands of mature proglotids (B). Detail of the prostatic glands is shown in C. *cy*: cytons, *ex*: excretory ducts, *p*: prostatic gland, *gp*: genital pore. Scale bars: 20 µm in A, 50 µm in B and 10 µm in C.
